# On the effect of inheritance of microbes in commensal microbiomes

**DOI:** 10.1101/2021.09.21.461237

**Authors:** Román Zapién-Campos, Florence Bansept, Michael Sieber, Arne Traulsen

## Abstract

**Background:** Our current view of nature depicts a world where macroorganisms dwell in a landscape full of microbes. Some of these microbes not only transit but establish themselves in or on hosts. Although hosts might be occupied by microbes for most of their lives, a microbe-free stage during their prenatal development seems to be the rule for many hosts. The questions of who the first colonizers of a newborn host are and to what extent these are obtained from the parents follow naturally.

**Results:** We have developed a mathematical model to study the effect of the transfer of microbes from parents to offspring. Even without selection, we observe that microbial inheritance is particularly effective in modifying the microbiome of hosts with a short lifespan or limited colonization from the environment, for example by favouring the acquisition of rare microbes.

**Conclusion:** By modelling the inheritance of commensal microbes to newborns, our results suggest that, in an eco-evolutionary context, the impact of microbial inheritance is of particular importance for some specific life histories.

## 1. Background

Microbial life is ubiquitous in the biosphere [1]. The human body is no exception, as first described by van Leeuwenhoek in the 17th century. We are among the many macroorganisms where diverse microbiomes – microbial communities living in or on hosts – have been observed [2, 3]. As part of their life cycle, members of the microbiome may migrate between hosts and the environment. The migration process has been studied using experimental [4] and theoretical approaches [5, 6]. However, some microbes have been found exclusively in hosts [4, 7]. How do such microbes persist in the population?

One possibility is the vertical transfer of microbes from parents to offspring [8]. Although there is ample literature about transmission of endosymbionts (e.g. *Buchnera* and *Wolbachia* in insects [9]), less is known about extracellular – possibly transient – microbes. Quantifying the low microbial loads in newborns [10] and deciphering the true origin of microbes [11] remains experimentally challenging [12, 13]. A few experimental studies have explored the vertical transfer of the microbiome in specific species across the tree of life – including sponges [14], mice [15], cockroach eggs [16], and wheat seedlings [17]. For many others, including humans, there is an ongoing debate on when and how inherited microbes are obtained [11]. Together, these studies suggest there is no universal reliance on microbial inheritance across host species, raising the possibility that even if such associations matter to the host, certain life-history traits may limit their inheritance [13, 18]. Relevant traits may include, among others, the extent of environmentally acquired microbes and host lifespan.

Previous theoretical work has studied microbial inheritance in the context of symbiosis – where microbes affect the host fitness. In these models, depending on whether the interaction is positive (mutualism) or negative (parasitism) the presence of symbionts is promoted or impeded, respectively. Using multilevel selection arguments, Van Vliet and Doebeli have shown that a symbiosis that is costly for microbes can be sustained only when the host generation time is short and the contribution of inheritance exceeds that of environmental immigration [19]. Following up, in addition to individual inheritance (single contributing parent), Roughgarden analyzed scenarios of collective inheritance (multiple contributing parents) [20]; while Leftwich et al. found a weak influence of the host reproductive mode (sexual or asexual) and mate choice (based on symbiont presence) on the symbiont occurrence [21]. If these host-symbiont interactions persist over evolutionary timescales, they are said to lead to phylosymbiosis – where microbiomes recapitulate the phylogeny of their hosts [22].

Not all co-occurrences between hosts and microbes reflect a fitness impact, however. As suggested by Bruijning et al., the selection on the host-microbiome pair and the microbial inheritance might change with the environment [18]. Moreover, despite taxonomic differences, functional equivalence of microbes in localized host populations could prevail [16]. Microbes might not always influence host fitness [18] nor benefit from influencing it [21]. In this context where there is no active selection of the microbes by the host, the role of microbial inheritance remains largely unexplored [23].

Using a stochastic model, we study the effect of microbial inheritance on the commensal microbiome – microbes living in hosts but not affecting their fitness. First, we introduce different models of inheritance representative of diverse host species. Then we discuss their effect on microbes present in both hosts and environment, or only present in hosts. We see that inheritance might influence the within-host occurrence and abundance in some cases. However, within the same microbiome, microbial types could be affected differently – while inheritance causes some microbes to increase in frequency, others decrease from it. Moreover, the effects may be transient, rendering life history parameters crucial. Altogether, we highlight the potential and limits of microbial inheritance to modify the composition of commensal microbiomes under different life-history scenarios.

## 2. Model and methods

Consider the host-microbiome system depicted in Fig. 1A. A population of hosts is colonized by a set of microbes, and each microbial taxon *i* has a constant frequency *p*_*i*_ in the environment. The total number of microbes a host can contain is finite and given by *N*. Each newborn empty host inherits a set of microbes from its parent, chosen at random within the host population. The inherited sample, taken off the parental microbiome, is drawn according to a probability distribution (Fig. 1B). After this initial seeding, only the death, immigration and replication of microbes can modify the host microbiome. Through these processes, the microbial populations within the host can decrease or increase by one individual each time step. After one microbe is selected to die, migration from the pool of colonizers occurs with probability *m*, while duplication of a resident microbe, or non-replacement, occurs with probability 1 − *m*. This process ends with the host death, which occurs with probability *τ* at each time step. We assume that the number of hosts does not change, so that a host death is followed by the birth of a new empty host, for which the process described above is repeated.

**Figure 1:**
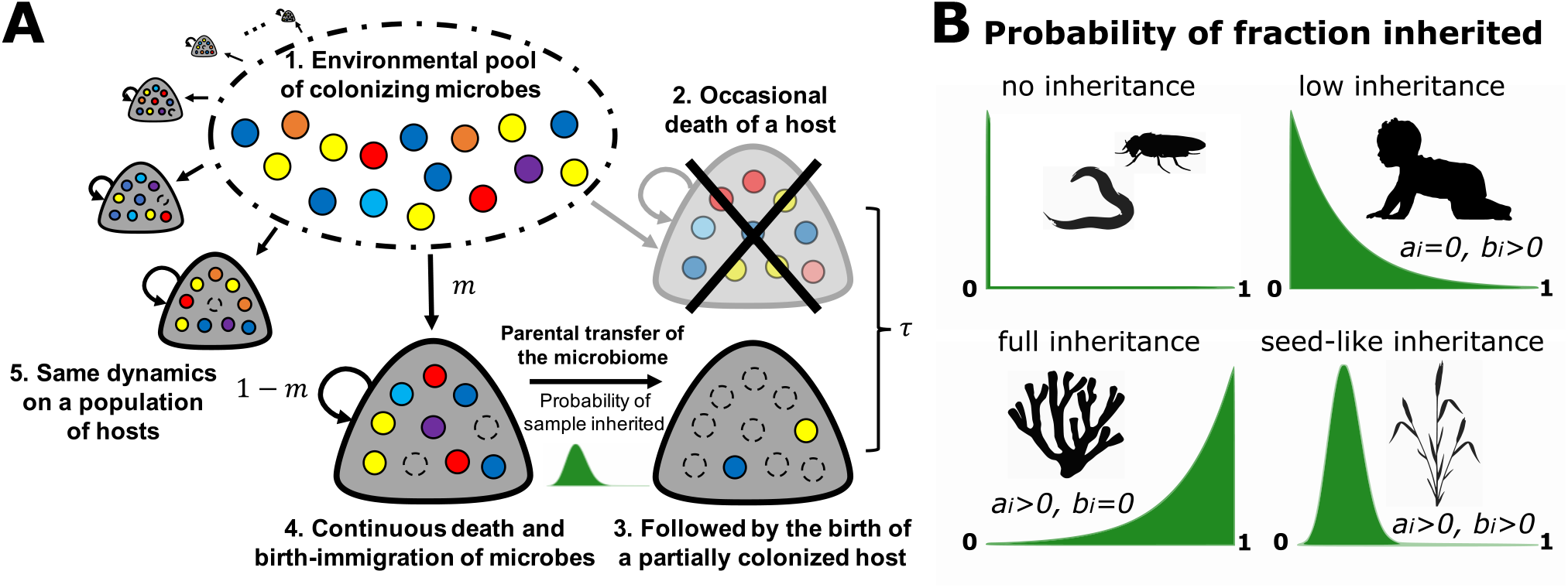
Host-microbiome dynamics and microbial inheritance in our model. (**A**) Dark blobs indicate hosts, coloured- and empty-circles indicate microbes and empty-space, respectively. Within the hosts, microbes go through a death and immigration-birth process, with new residents migrating from the pool of colonizing microbes with probability *m* or replicating within a host with probability 1 − *m*. For microbes, each host is an identical habitat. The host population is at a dynamic equilibrium, every timestep there is a probability *τ* that a host death occurs, immediately followed by the birth of a new one. The newborn obtains a sample of its parent microbiome according to a probability distribution. (**B**) The probability distribution of the fraction of the parental microbiome inherited vary across host taxa – among others, influenced by development, reproduction and delivery mode. Certain hosts might not transfer microbes (eg. *C. elegans* [24] or *D. melanogaster* [25]). Others might provide minimal (eg. humans [11]) or large fractions of their microbes (eg. fragmentation of some sponges, corals, fungi and plants [26, 27]), while others might be centred around a fixed value (eg. seeds of plants [17]). In our model, we control this probability distribution through the parameters *a*_*i*_ and *b*_*i*_ in Eq. (4).

### 2.1. Transition probabilities

Our aim is to describe the dynamics of the microbiome load and composition, focusing in particular on how a certain microbial taxon experiences it. Within a specific host, the frequency of the *i-th* taxon is denoted by *x*_*i*_ (for *i* ≥ 1), and of the remaining other microbes by *o*_*i*_ = Σ_*j ≠ i*_ *x*_*j*_. The frequency of available space is then given by *x*_0_ = 1 − *x*_*i*_ − *o*_*i*_. The transition probabilities from state {*x*_*i*_, *o*_*i*_} that are due to the microbial dynamics are given by the product of the probability of host survival, 1 − *τ*, by the probability of death of a certain microbial type followed by an immigration or birth event. These events produce changes in the frequencies of magnitude 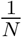. First, microbial taxa can replace each other when a microbe dies and is replaced by another one,

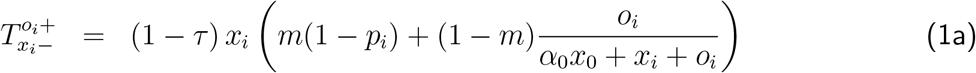

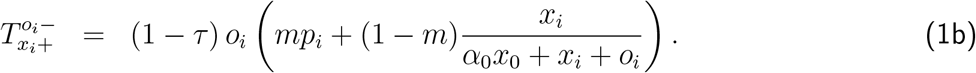

In Eq. (1a), a microbe of type *i* dies and is replaced by another microbe, either by immigration from the environmental pool or by replication within the same host. Similarly, in Eq. (1b), a microbe of another type dies and is replaced by a microbe of type *i*.

Alternatively, microbes may occupy previously available space, such that the microbial abundance increases,

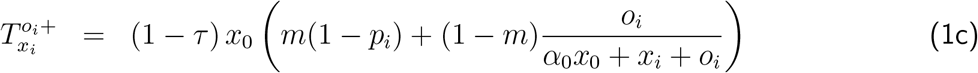

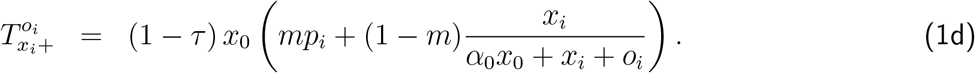

Finally, microbes may decrease in abundance, when a microbe selected for death is not replaced,

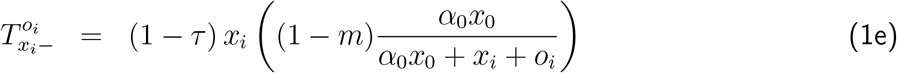

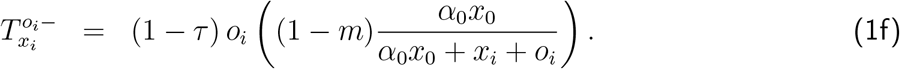

In these equations, *α*_0_ controls the establishment of microbes in hosts – the ability to occupy available space – going from fast for *α*_0_ = 0, to slow if *α*_0_ is positive. For *α*_0_ > 1 and without migration, microbes cannot be maintained in hosts.

The transition probabilities due to the hosts dynamics are given by the product of the probability of host death and birth of an empty host (*τ*), by the probability to inherit certain microbes,

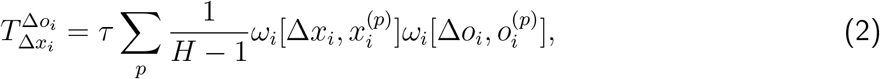

where 1*/*(*H* − 1) is the probability of choosing a parent *p* in the population of *H* − 1 potential parents, and 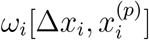 and 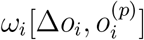 are the probabilities of transfer of Δ*x*_*i*_ and Δ*o*_*i*_ microbes from the parent to the offspring, respectively. Because the frequencies within the parent are 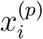 and 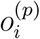, the probability to transfer more microbes than the parent can provide is zero.

Finally, for completeness, the probability of staying in state {*x*_*i*_, *o*_*i*_} without host death is

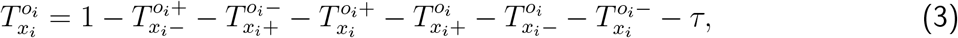

where the last term includes all possible transitions due to parental transfer of microbes, 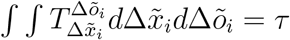.

### 2.2. Distribution of inherited microbes

In our model, parents can seed the microbiome of their offspring. A sample of the parental microbiome is vertically transmitted according to a probability distribution function, Eq. (2). In addition to the case without inheritance, which we have previously analyzed elsewhere [28], at least three qualitatively distinct cases may be observed (Fig. 1B), depending on host development, reproduction, and mode of delivery.

Firstly, inheritance could be low. For example in animals, newborns get microbes attached to epithelia or fluids during delivery [11, 8]. These represent a small fraction of the parental microbiome, leading to distributions centred at frequency zero decaying towards one. Secondly, certain hosts, including some sponges, corals, fungi and plants [26, 27], are able to reproduce by fragmentation, where a breaking body part generates a new individual. Such fragments could carry a faithful microbiome composition, leading to distributions centered at frequency one decaying towards zero. Finally, hosts that produce embryos that can disperse, eg. seeds, might transfer a microbiome sample contained within these physical structures [17].

We modelled such diverse parental microbiome samplings (Δ*x*_*i*_) using the beta distribution for the probability 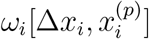 to inherit Δ*x*_*i*_ microbes from parent *p*. This probability distribution can take arguments in the range from zero to the current frequency of a microbe in the parent *p*, 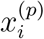,

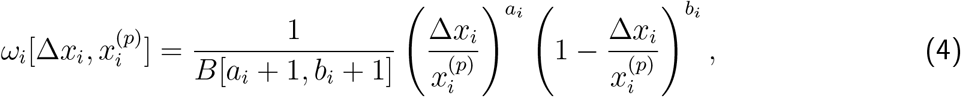

where *B* is the beta function [29], 1*/B* a normalization constant, and *a*_*i*_ and *b*_*i*_ are shape parameters. The expected value of our beta distribution is 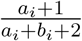. The special case of *a*_*i*_, *b*_*i*_ = 0 leads to a uniform distribution, where the parental microbes are distributed randomly between parent and offspring. Other combinations of *a*_*i*_, *b*_*i*_ ≥ 0 produce different unimodal distributions (Fig. 1B). The case of *a*_*i*_ *> b*_*i*_ skews the distribution towards full inheritance of the parental microbes, 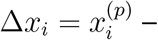 all the *i-th* microbes from the parent could be transferred to the offspring. The case of *a*_*i*_ *< b*_*i*_ skews the distribution towards non-inheritance of microbes of type *i* to offspring, Δ*x*_*i*_ = 0. Finally for *a*_*i*_ = *b*_*i*_, the distribution is symmetric and the parental microbes are likely to be equally distributed between parent and offspring. In most of our analyses *a*_*i*_ and *b*_*i*_ are the same for all microbial taxa. Only for non-neutral, asymmetric inheritance, we will set different *a*_*i*_ and *b*_*i*_ for the focal taxon (*x*_*i*_) and the set of others (*o*_*i*_). To illustrate the effect of *a*_*i*_ and *b*_*i*_, on average, an offspring inherits *≈* 9% of the parental microbes of taxon 1 for *a*_1_ = 0 and *b*_1_ = 9, while only *≈* 1% is inherited for *a*_1_ = 0 and *b*_1_ = 99.

Throughout the results, we focus on distributions with a maximum at microbial frequency zero decaying towards 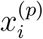, which we call ‘low inheritance’ (Fig. 1B). In our model, the low inheritance and the ‘full inheritance’ scenarios (distributions with maximum at frequency 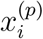 decaying towards zero) are equivalent. This stems from the fact that the number of microbes is conserved, so that inheritance happens through the splitting of the parental microbiome between the parent and the offspring. Thus, since in our model, the probability to die of a host does not depend on its age, the splitting of microbes in the low inheritance scenario - where a small fraction is transmitted - and in the full inheritance scenario - where most of the microbiome is transmitted - are equivalent. Finally, we address under which circumstance a ‘seed-like inheritance’ leads to different results.

### 2.3. Stochastic simulations

In order to simulate the microbiome dynamics of individual hosts we formulated the model as a stochastic differential equation. We solved this equation numerically using the Euler-Maruyama method [30]. Starting from state **x** = {*x*_*i*_, *o*_*i*_} at time *t* the new state after an interval Δ*t* is given by

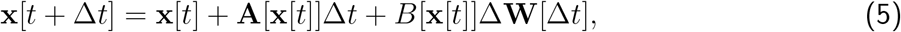

where **A**[**x**[*t*]] is the vector of expected changes of **x**, the deterministic contribution; while *B*[**x**[*t*]] is a matrix that has the property *B*[**x**[*t*]]^*T*^ *B*[**x**[*t*]] = *V* [**x**[*t*]], where *V* [**x**[*t*]] is the covariance matrix of the change of **x**. Further, Δ**W** is a vector of uncorrelated random variables sampled from a normal distribution with mean 0 and variance Δ*t*, the stochastic contribution. That Δ**W** is normally distributed arises from the time independence and identical distribution of the noise. A detailed description connecting Eq. (1) and Eq. (5) is provided in Appendix A1.

For most of their life, hosts are independent of each other, only newborns are influenced by others when they acquire their initial microbiome. A given host lives for a duration sampled from an exponential distribution *τe*^−*τt*^, with mean 1*/τ*. We solve Eq. (5) for that interval. Immediately after a host dies, the microbiome of a newborn is assembled according to Eq. (2). We repeat these steps for all hosts until the total simulation time is reached.

As a result of stochasticity, each host trajectory is different. We look into the statistical description of the microbiome composition across the host population.

## 3. Results

### 3.1. Inheritance can increase the occurrence of microbes in hosts with low microbial loads

Without microbial inheritance, which will be our reference case throughout, any microbe occurring inside a host has to have migrated from the environment during the host lifespan. As a result, a low environmental migration or short host lifespan can be limiting [28]. The transfer of microbes from parents to offspring during birth could increase the probability of observing any microbes in hosts, *P* [*x*_*i*_ + *o*_*i*_ > 0]_inh._. We quantified the change in the probability of occurrence relative to its microbe-free birth condition *P* [*x*_*i*_ + *o*_*i*_ > 0]_no inh._,

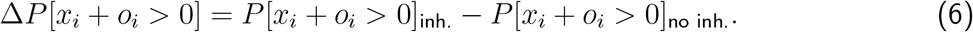

Using this observable, we investigated the role of life history in modulating the effect that inheritance has on the microbiome. We quantified this for a single microbial taxon, *x*_*i*_, as well.

Fig. 2 shows a condition where, in the absence of inheritance, hosts are not fully occupied by microbes. This results from a short host lifespan (*τ*) and low microbial immigration from the pool of colonizers (*m*). We tested the effect of the ‘low inheritance’ mode (Fig. 1B) for increasing rates of establishment of microbes (*α*_0_ → 0) and other life-history parameters.

**Figure 2:**
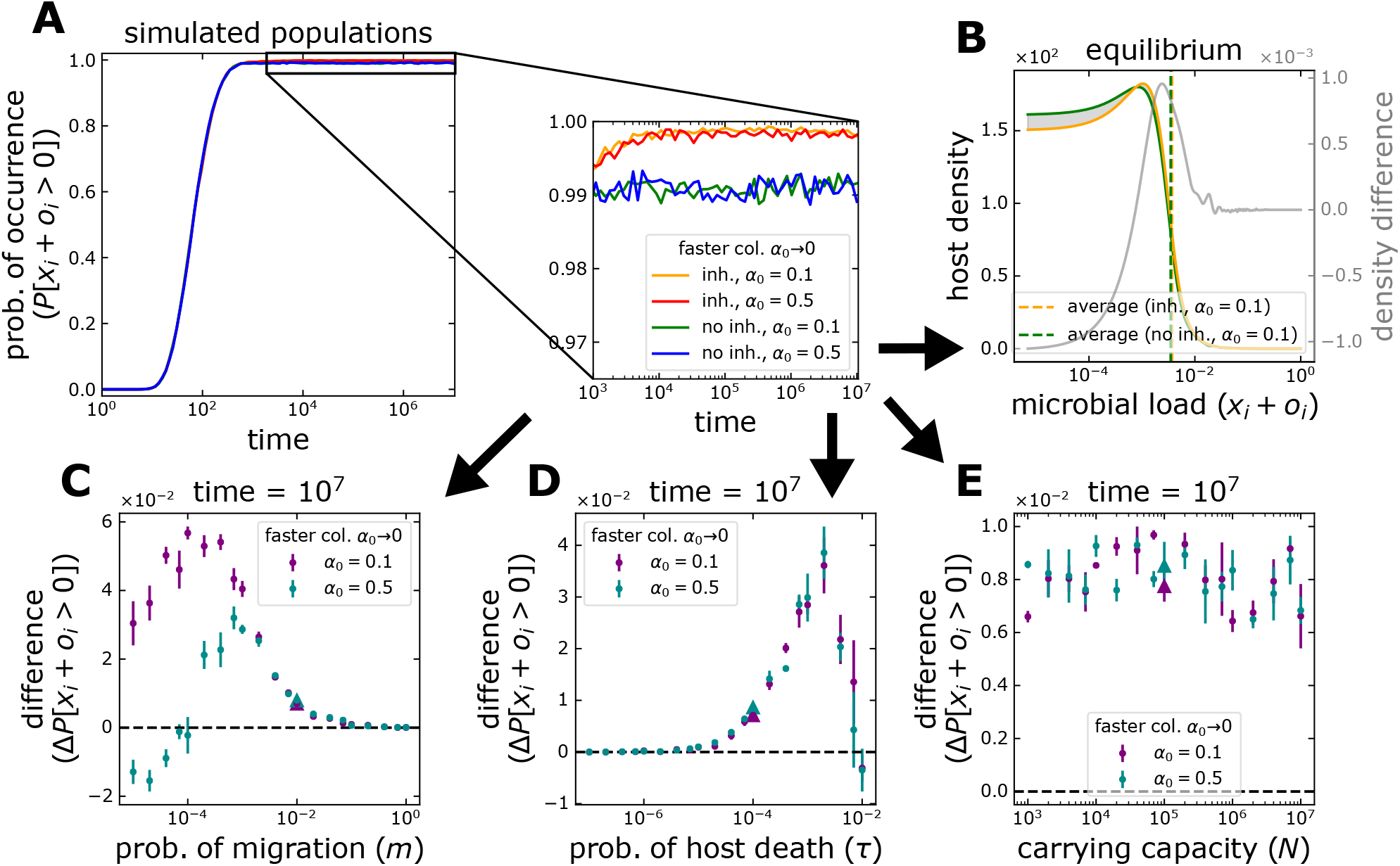
Microbial occurrence in hosts under microbial inheritance. (**A**) Starting from a condition where all hosts are initially empty, the microbial occurrence increases through time. At first sight, this increase is largely independent of *α*_0_ and the inheritance of microbes. A closer look at equilibrium abundance reveals that inheritance increases the occurrence, in this case, regardless of how rapidly hosts are occupied (*α*_0_). (**B**) The increase results from a distribution of microbial load across the host population where the microbe-free state is less common. A microbial load of 10^−5^ corresponds to 1 microbe per host. In (C-E), single parameters are modified from the case shown in (A-B) (with parameters *m* = 10^−2^, *τ* = 10^−4^, and *N* = 10^5^, indicated by the triangles in (C-E)). (**C**) A large migration from the pool of colonizers, *m* → 1, hinders any effect of inheritance on occurrence as hosts are readily colonized. The change peaks and decreases for smaller *m*, as for *m* → 0 hosts are less likely to be colonized. The change can even be negative for slowly occupied hosts where the few colonizing microbes are lost to stochasticity. (**D**) The gain from inheritance is maximal for intermediate values of host death probability, *τ*. Long living hosts, *τ* → 0, are colonized even without inheritance. Short living hosts, *τ* → 1, are less likely to be colonized and thus transmit microbes through inheritance. (**E**) The carrying capacity for microbes of a host, *N*, and *α*_0_ do not alter the gain from inheritance. Points and bars in (C-E) indicate the average and standard deviation of 6 simulation pairs, with vs. without inheritance, with 10^4^ hosts each. Offspring receive 9% of their parent’s microbiome on average, *a*_*i*_ = 0 and *b*_*i*_ = 9 in Eq. (4). The whole distributions are shown in Fig. Sup. 2.

Inheritance impacts the occurrence of microbes by increasing the number of hosts with at least one colonizing microbe (Fig. 2B). The effect is most prominent in scenarios where without inheritance, most of the hosts are microbe-free. However, the maximum increase occurs at intermediate immigration and host lifespans (Fig. 2C-D). For high immigration, *m* → 1, hosts are readily occupied by microbes, so inheritance brings no change. Similarly for a long host lifespan, *τ* → 0. On the other hand, if immigration is limited, *m* → 0, or host lifespan short, *τ* → 1, microbes never occur in hosts, so parents cannot transmit microbes to their offspring.

Inheritance might decrease the occurrence if the transfer – which splits the parental microbiome between parent and offspring – makes microbes more susceptible to stochastic fluctuations. This occurs if the microbial frequency of the parent is already low – for example when migration is limiting and microbes proliferate slowly (Fig. 2C). This phenomenon might be pronounced for individual taxa. Our analyses from the perspective of a single taxon (Fig. Sup. 1) found multiple instances where inheritance might decrease the occurrence (Fig. Sup. 1C-F), but also have a larger effect in situations where the occurrence increases. Additionally, the effect on single taxa depends strongly on the carrying capacity for microbes, *N* (Fig. Sup. 1F compared to Fig. 2E). Competition for space favours taxa according to their frequency in the pool of colonizers, *p*_*i*_ (Fig. Sup. 1C). Abundant taxa outcompete rare ones as space is more limited, but only until a point, after which there is no benefit – they readily occur without inheritance. In other words, in microbiomes composed by many taxa, the taxon-level effect of inheritance in terms of occurrence is relative to their environmental abundance.

### 3.2. Inheritance can increase the abundance in hosts, but mostly of those abundant in the environment

Modifying the presence of taxa is not the only effect – inheritance also alters the microbiome composition considerably. Using the distribution of microbial frequencies in hosts, we quantified the change in the average frequencies as compared to its microbe-free birth condition,

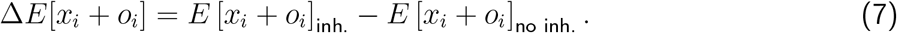

Similarly to Eq. (6), we quantified this observable for a single microbial taxon, *x*_*i*_, as well.

When looking at the distribution of microbial loads and frequencies in hosts, the effect of the ‘low inheritance’ mode (Fig. 1B) is two fold – while hosts with small frequencies might experience the largest increase in microbes, hosts with large frequencies can see the largest decrease of microbes (Fig. 2B and Fig. Sup. 2). Thus, at both microbial load and single taxon levels, hosts with small and large frequencies become rarer. Inheritance makes hosts resemble each other to a greater extend (see the reduced spread of the distributions in Fig. Sup. 2 and Fig. Sup. 3). This is equivalent to the effect of increased immigration, which also tends to make microbiomes similar to one another, but increased inheritance does not favour the preservation of the diversity from the pool of colonizers – in contrast to immigration.

An increase in the average load is observed for some conditions (Fig. Sup. 2). Analogously to the occurrence, such increase peaks at intermediate host death probabilities *τ*; but also at intermediate carrying capacities *N* (Fig. 3C-D). The limited time for host colonization impedes any inheritance (*τ* → 1), while for *τ* → 0 or small *N*, hosts are fully occupied even without it. The relative effect of inheritance is less for large *N*. A faster occupation of available space (*α*_0_ → 0) displaces the effect to larger host death probabilities and capacities for microbes. Finally, because the main limitation is the short host lifespan (*τ*), the influence of immigration (*m*) is minimal (see the scale in Fig. 3B and Fig. Sup. 4C).

**Figure 3:**
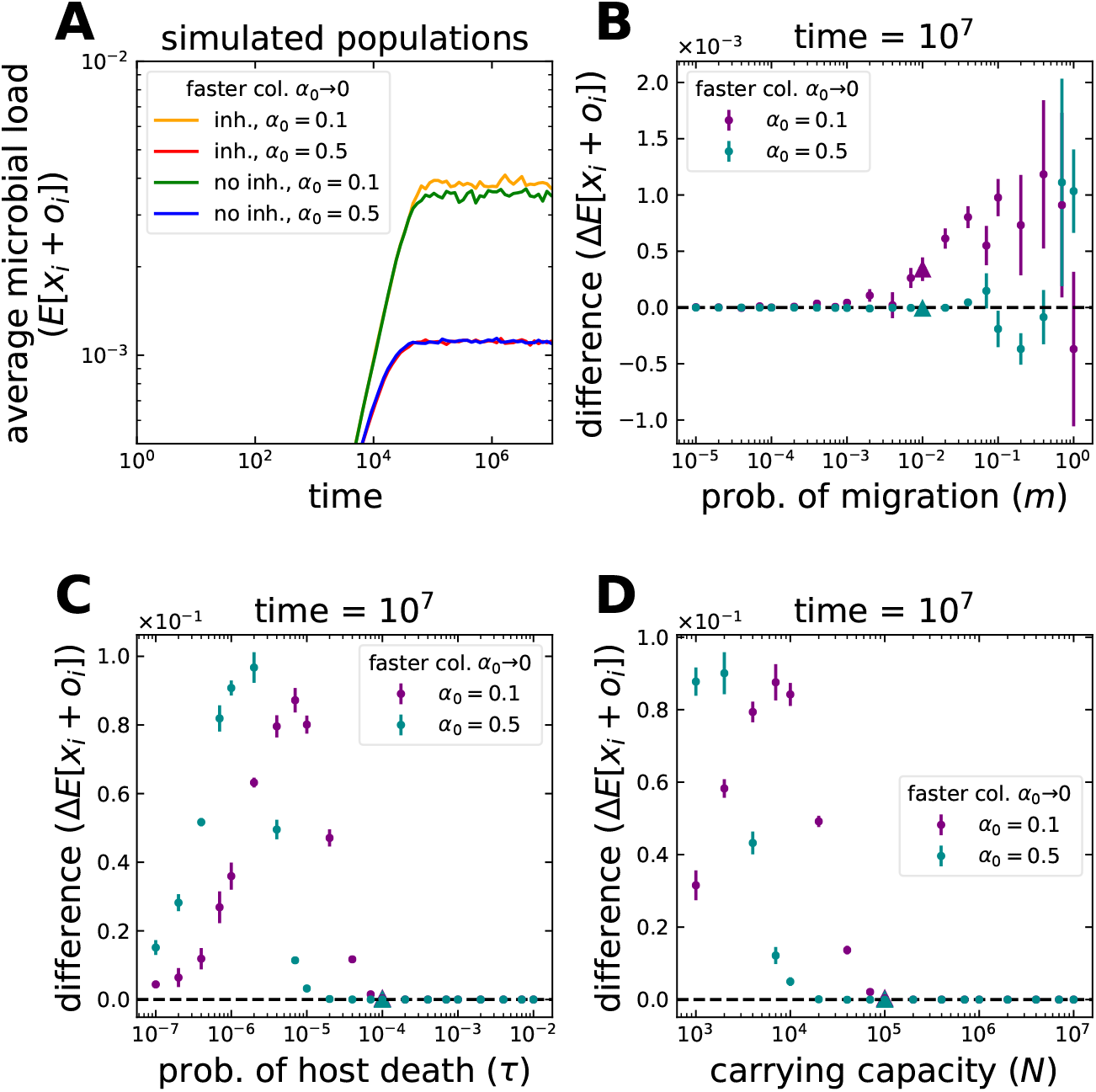
Average microbial load in hosts under microbial inheritance. (**A**) Starting from a condition where all hosts are initially empty, the average frequency of microbes in hosts increases through time before reaching an equilibrium. In this particular case, inheritance makes such equilibrium abundance larger only when hosts are occupied rapidly, *α*_0_ → 0. This increase results from a host distribution where higher microbial loads are more common (Fig. 2B). The cases shown in (A), with parameters *m* = 10^−2^, *τ* = 10^−4^, and *N* = 10^5^, are indicated by the triangles in (B-D). A single parameter is varying in (B-D). (**B**) Changes of migration from the pool of colonizers, *m*, have minimal effect (notice the scale). As *m*→ 1, more microbes colonize the hosts. Still the average microbial load only increases if the loss of microbes to inheritance is less than the gain from proliferation. (**C**) The effect of changes to host death probability, *τ*, are much larger and maximal at intermediate *τ*. A faster occupation of hosts makes the effect of inheritance larger for shorter living hosts, *τ* → 1. (**D**) In contrast to the occurrence (Fig. 2E), changes in the carrying capacity for microbes, *N*, have a larger intermediate effect. Faster occupation of hosts makes the effect peak for larger *N*. Points and bars in (B-D) indicate the average and standard deviation of 6 simulation pairs, with vs. without inheritance, with 10^4^ hosts each. Offspring receive 9% of their parent’s microbiome on average, *a*_*i*_ = 0 and *b*_*i*_ = 9 in Eq. (4). The whole distributions are shown in Fig. Sup. 2.

Although higher loads might be reached with inheritance if space is limited (Fig. Sup. 2C), abundant taxa might increase at the expense of rare ones (Fig. Sup. 3D and Fig. Sup. 4D-E). Such reduction is exacerbated by the fast occupation of available space *α*_0_ → 0. Interestingly, this might happen as a result of longer host lifespans as well, if hosts are rapidly occupied by inherited microbes. Such condition favours abundant taxa in the pool of colonizers. Instead, if the occupation is slower, rare taxa increase in frequency, derived from the added benefits of inheritance and a more influential immigration (*m*).

A particularly relevant question is whether the frequency of a taxon in a specific host (*x*_*i*_) can be larger than in the pool of colonizers (*p*_*i*_) – i.e. a benefit is obtained from the host association. We observe this even in the absence of inheritance (Fig. Sup. 3), where stochastic colonization results in some host containing frequencies larger than in the pool (*p*_*i*_). The average frequency across hosts, however, can be larger only when the space limitation increases the competition. In this context, inheritance may, in fact, decrease the chances of such outcome, by relating the hosts to each other (Fig. Sup. 3C-D).

### 3.3. Preferential inheritance can temporally lead to specific taxa overrepresentation

A potential mechanism to increase the average frequency of taxa beyond their frequency in the pool of colonizers (*p*_*i*_), is preferential inheritance. The asymmetry in inheritance could stem from differences in microbial properties, but also a host’s direct or indirect influence. We studied such possibility by manipulating the distribution of the sample inherited, Eq. (4). Focusing on a ‘low inheritance’ mode, we decreased the inheritance of other taxa relative to taxon *i*, from equal if offspring receive 9% of every taxa on average, to preferential if they receive 9% of taxon *i* but 1% of others.

For the same parameters as before (Fig. 4), we observe no effect if the host lifespan is limiting. In this case, regardless of the frequency in the pool of colonizers (*p*_*i*_), preferential inheritance does not alter the average frequency of the *i-th* taxon in hosts (Fig. 4A), similarly for the probability of immigration *m* (Fig. 4B). This holds even for fast occupation of available space, *α*_0_ → 0. Only for longer host lifespan, *τ* → 0, preferential inheritance leads to an increase (Fig. 4C). Besides the almost exclusive occupation of hosts by the *i-th* taxon 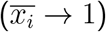, the maximum effect is constrained to intermediate *τ*. This is because the effect of preferential inheritance is transitory for longer living hosts, after which they continue approaching their long term equilibrium, 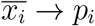. For faster occupation of available space the gain spans a wider range and shorter host lifespans (*τ* → 1). For hosts with short lifespan and limited immigration (in our example *τ* = 10^−4^ and *m* = 10^−2^), the gain from preferential inheritance is largest for decreasing carrying capacity for microbes, *N* (Fig. 4D). As shown in Fig. 4D, inheritance itself might not benefit all microbial taxa. For some taxa, only preferential inheritance can lead to larger frequencies than without inheritance.

**Figure 4:**
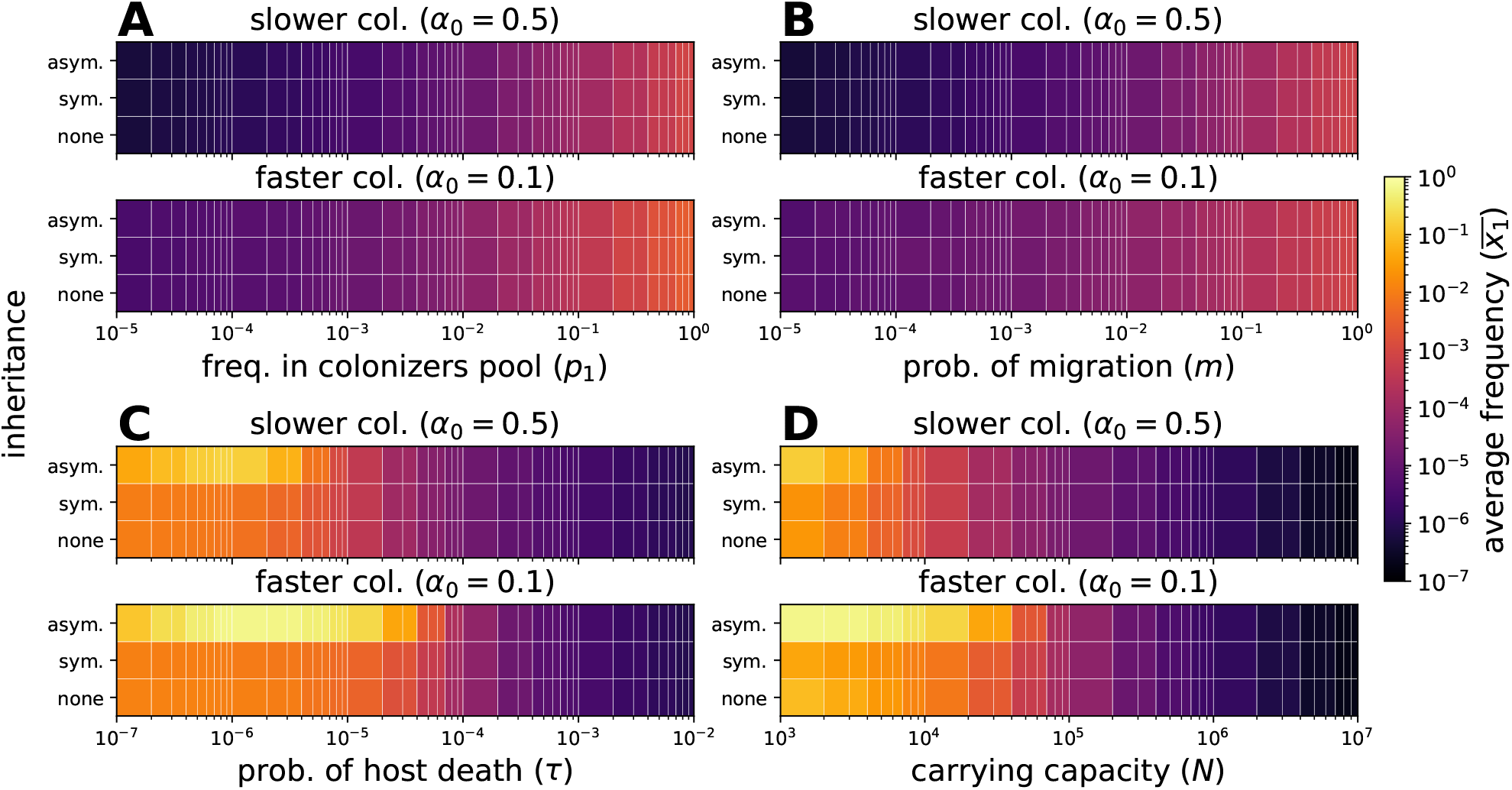
Effect of asymmetric inheritance on the average frequency of a taxon in hosts. Cases without inheritance and inheritance are compared. Inheritance is symmetric if offspring receive 9% of their parent’s microbiome on average (*a*_*i*_ = 0 and *b*_*i*_ = 9). Inheritance is asymmetric if offspring receive 9% of taxon 1 and 1% of other taxa (*a*_*i*_ = 0 and *b*_1_ = 9, *b*_*i*≠1_ = 99 in Eq. (4)). Available space within hosts is occupied more easily for *α*_0_ → 0. Single parameters are modified from the condition *p*_1_ = 10^−2^, *m* = 10^−2^, *τ* = 10^−4^, and *N* = 10^5^. (**A-B**) The average frequency increases for larger abundances in the pool of colonizers (*p*_1_), immigration (*m*), and *α*_0_ → 0. An asymmetric inheritance has no effect, as hosts are not fully occupied within their lifetime (Fig. Sup. 2 and Fig. Sup. 3). (**C**) Longer host lifespans, *τ* → 0, increase the average frequency and effect of asymmetric inheritance. The gain is maximal at intermediate *τ*. Inheritance has more influence before hosts are fully occupied. After this, hosts resemble the colonizers pool. (**D**) The average frequency increases with competition for space (smaller *N*). While the symmetry of inheritance decreases the average frequency as a result of the reduced initial microbiome variability, asymmetry increases it. Each simulation included 10^4^ hosts.

### 3.4. Persistence of lineage taxa in hosts

An extreme case of reliance on microbial inheritance are microbes present in hosts but absent from the environment (*p*_*i*_ = 0) [1, 7]. We refer to these as lineage taxa. We investigated the conditions allowing their persistence under different life-history scenarios (Fig. 5).

**Figure 5:**
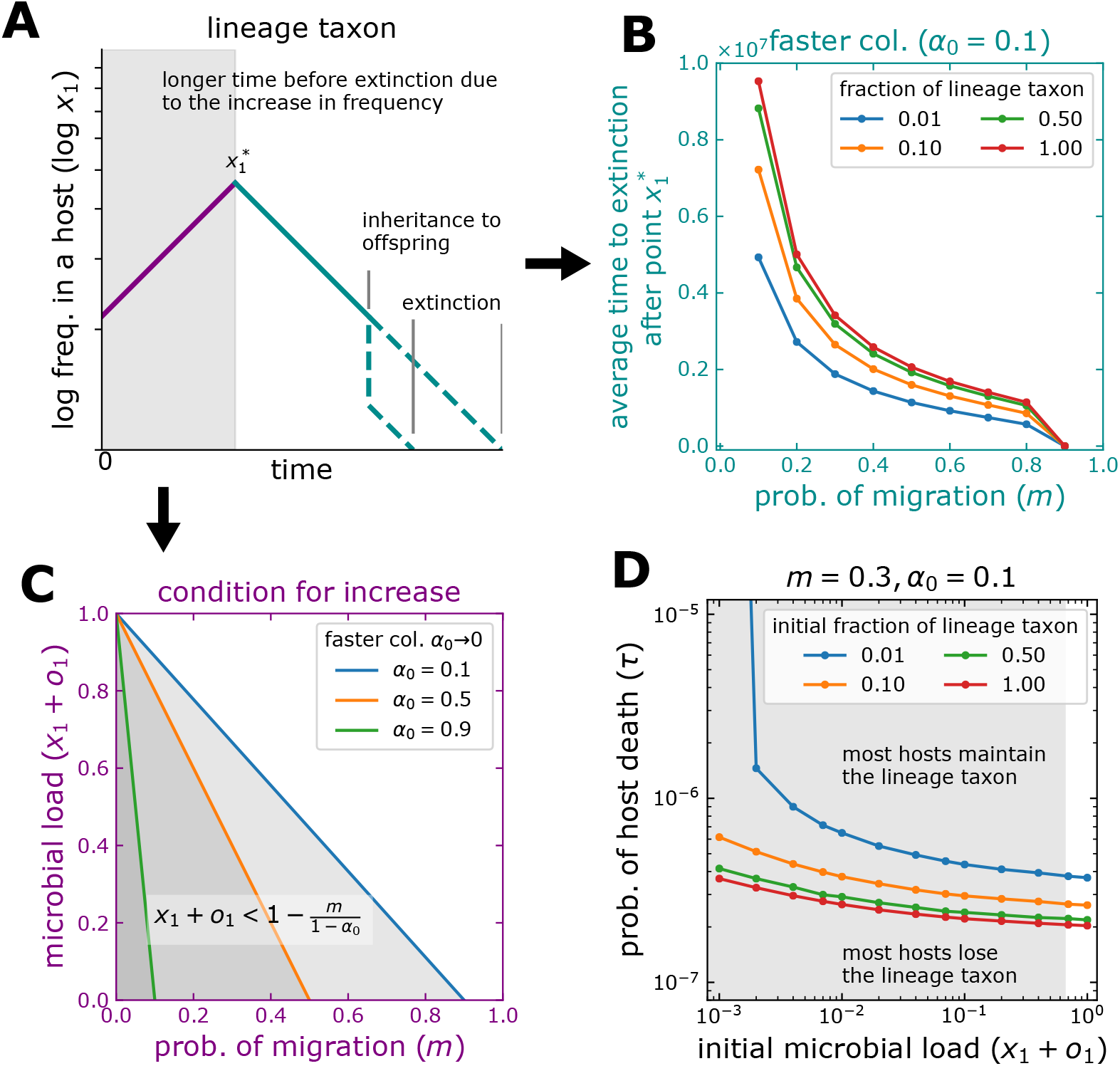
Persistence of lineage taxa in hosts. A microbial taxon is initially present in hosts *x*_1_(0) > 0, but not in the pool of colonizers, *p*_1_ = 0. (**A**) The frequency within a host decreases through time. For some conditions, Eq. (8), there is a period of increase. If the taxon is transmitted to offspring before the gain is lost, this might persist in the host population (although extinction within the parent occurs sooner). (**C**) Low immigration (*m* → 0) and fast occupation of available space (*α*_0_ → 0) allow increase and prolong the time before extinction, Eq. (8). Large initial available space (*x*_*i*_+*o*_*i*_ → 0) and lineage taxon fractions (*x*_1_*/*(*x*_1_ + *o*_1_) → 1) also prolong this time. (**B**) After the increase stops 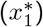, the average time to extinction is shorter for large immigration (*m* → 1) and a smaller fraction of the taxon. (**D**) At the host population level, lines indicate the death probability after which most hosts lose the lineage taxon (*τ*_0.5_), Eq. (9). The early increase shown in (A) only occurs within the darkened area. The distribution of microbes inherited, Fig. 1B and Eq. (4), affects the initial load and fraction of lineage taxa in offspring. Asymmetric inheritance in low microbial loads might preserve lineage taxa as well as symmetric inheritance in high loads. We set *N* = 10^5^. Each point corresponds to 10^4^ simulated hosts.

Within a host, lineage taxa go through the stages sketched in Fig. 5A. Depending on the context, after host birth, their frequency might either decrease or increase. If decrease occurs, in a neutral context this trend will not change during the host life. In fact, events of microbiome inheritance will further decrease the frequency in the parent. We found that on average, lineage taxa increase while the inequality

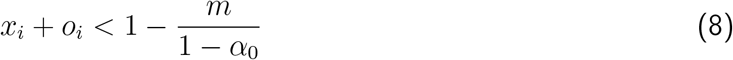

holds (Fig. 5C and Appendix A). Therefore, lineage taxa increase before reaching carrying capacity, favoured by their fast proliferation (*α*_0_ → 1), but restricted by migration (*m*). Because the microbial load increases through time (*x*_*i*_ + *o*_*i*_ → 1), alongside the initial state, Eq. (8) limits the time of increase. Note that on average, the maximum frequency of lineage taxa is 1 − *m/*(1 − *α*_0_). From this point on, a decrease driven by the immigration of environmentally present microbes (*m*) and stochasticity follows. For sufficiently long time, such decrease may lead to their extinction (Fig. 5B). There is a trade-off between the duration of the increase and the maximum frequency of lineage taxa. While small initial microbial loads lead to long durations but small frequencies (as a result of immigration, Eq. 8), the opposite is true for high initial loads abundant in lineage taxa. Once increase stops, the time to extinction is proportional to the lineage taxa frequency, Fig. 5B. Putting these two times together, the extra time from the increase is behind the subtle effect of the initial microbial load on the total extinction time, Fig. 5D. A reduced migration (*m* → 0) and fast occupation of available space (*α*_0_ → 0) simultaneously increase the frequency and time before extinction.

Looking at the population level, a condition for persistence emerges – namely, an increase of frequency in each host followed by transfer to offspring of a frequency at least equal to that received at birth. This is possible only while the frequency in the parent is larger than initially, Fig. 5A. The largest frequencies are expected at intermediate time. In this context, host lifespan, and thereafter the probability of host death (*τ*) become fundamental. From the distribution of host death events, *τe*^−*τt*^, we see most hosts die early on, potentially while lineage taxa are still abundant; *τ* → 0 results in longer living hosts – those more likely to lose lineage taxa. We estimated the probability of host death at which a fraction *z* of hosts loses the taxa,

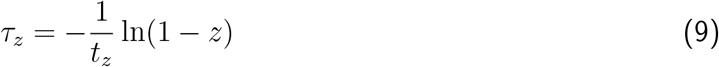

where *t*_*z*_, the time at which lineage taxa remain present in a fraction *z* of the host population, is obtained from the distribution of extinction times. Based on the former observations (Fig. 5D), our model predicts that regardless of the distribution of inherited microbes (Fig. 1B), preferential inheritance of lineage taxa in small microbial loads might favour their persistence as effectively as large but non-preferential microbial loads.

### 3.5. When the distribution of inherited microbes matters

We proposed that a finite set of shapes captures most of the possible microbial inheritance distributions (Fig. 1B) – low, high, and seed-like inheritance – all characterized by the most likely microbiome fraction transferred to the offspring. So far, we have focused on the impact of low inheritance on the microbiome (Fig. 2-5). As mentioned before, because we enforce the conservation of microbes in our model, i.e. the microbes are transferred from the parent host to the offspring, the outcome of low and high inheritance is equivalent: although the parental microbiome is distributed differently, the outcome is indistinguishable at the host population level, because hosts are indistinguishable.

When referring to certain life-histories, other distribution shapes may alter the impact of inheritance. To find out differences between the effect seed-like inheritance and our former results (where we assumed low inheritance) we compared the occurrence and average microbial frequencies.

We found most changes are minimal, however, differences appear for extreme parameters. A seed-like inheritance might better guarantee the occurrence of microbes in extremely adverse life-histories – e.g. rare environmental migration (*m* → 0) and short host lifespan (*τ* → 1) simultaneously (vertical axis on Fig. Sup. 5A-B). Exceptions could arise for a slower occupation of available space (*α*_0_). For individual microbial taxa, changes are greater in occurrence as well (Fig. Sup. 6); however, derived from the competition for limited space (*N*), the effect of a seed-like inheritance is case-specific. Moreover, both maximum increase and maximum decrease occur at intermediate *m* (Fig. Sup. 6B) and *τ* (Fig. Sup. 6C). In microbiomes composed of taxa with different environmental frequencies (*p*_*i*_), while some taxa gain, others lose from inheritance (Fig. Sup. 6A).

Under less adverse conditions, seed-like inheritance might allow larger microbial loads. That is the case when either host lifespan (horizontal axis on Fig. Sup. 5A) or migration (Fig. Sup. 5B) is limiting. The gain from a seed-like inheritance can be large, especially for a small carrying capacity for microbes *N* (Fig. Sup. 5C). The consistent microbial transfer and reduced variation are beneficial. Nonetheless, at the single taxon level, gains are minimal (Fig. Sup. 6). At this level, a limiting carrying capacity for microbes, where competition increases, might even lead to a decrease (Fig. Sup. 6D). In this case, the variation provided by the low inheritance mode is more beneficial.

In summary, regardless of the distribution of microbes inherited (Fig. 1B), life-history seems intrinsically linked to the effect of microbial inheritance on the microbiome composition.

## 4. Discussion

The impact of microbial inheritance on host-associated microbial communities is largely unknown. In this work, we explored its potential effects under diverse life-history scenarios, including multiple distributions of microbes inherited (Fig. 1). Using a model free of selection – i.e. without microbial fitness differences or effect on host fitness – we shed light on the conditions where microbial inheritance may influence the microbiome composition, showing its impact but also its limits.

Our work emphasizes the role of life-history over host-microbe associations (Fig. 2-3). Even without symbiotic benefits, the inheritance process itself might alter the microbiome composition [21]. Using a discrete generation model, Zeng et al. considered microbial inheritance in neutral associations over evolutionary timescales – specifically, its effect on the microbial diversity and the distribution of frequencies [23]. Our results, however, highlight the relevance of within-generation probabilistic events – environmental colonization, host lifespan, or carrying capacity for microbes – as ecological drivers to constrain inheritance.

A crucial constraint is the host lifespan. Similarly to Van Vliet and Doebeli, but without any impact on the host fitness, we observe that the environmental acquisition of microbes makes the effects of inheritance transient (Fig. 2D, 3C and 4C) [19]. Short-living hosts (relative to the microbial timescale) could influence their commensal microbiome over their whole lives, while long-living hosts only during the first stages of development. The rapid proliferation of inherited microbes or isolation from the environment might prolong the period of influence. This is in contrast to Van Vliet and Doebeli, where selection within isolated hosts acts against costly symbiosis, reducing the period of mutualists presence.

We observed that the effect of inheritance may differ between taxa. Microbiomes assembled entirely from the environment are prone to variation when migration between hosts is rare [28, 18]. Inheritance might increase the presence of certain microbes, but in contrast to environmental migration, inheritance reduces the variation between hosts and potentially their microbial diversity. This reduction, which especially affects rare taxa, is more pronounced if the carrying capacity is limited (Fig. Sup. 1 and Sup. 4), where competition is larger. Bruijning et al. have observed that under selection, such decreased variation and diversity could be detrimental for adaptation to changing environments [18].

Initially, we assumed no distinction between microbial taxa – only their frequency determined the population dynamics (Eq. 1). This could be modified in at least two ways. First, fitness differences could influence the birth and death rates of microbes. Although this is certainly relevant, it diverts from our focus on inheritance. Instead, we addressed a possibility crucial for inheritance – the asymmetric transfer of microbes (Fig. 4). Such asymmetry could emerge from differences in microbial capabilities at play during the transfer process, including oxygen tolerance [15] (obligate anaerobes tend to be transmitted vertically) and sporulation [31] (spores might allow the transfer of oxygen-sensitive bacteria). Alternatively, hosts could selectively transfer certain microbes to their offspring [9]. Interestingly, we observe that inheritance alone is not always beneficial; some taxa might only benefit when transferred asymmetrically (Fig. 4).

We have emphasized the importance of looking at rare taxa. Such is the case of lineage taxa (Fig. 5), microbes absent from the environment that only propagate by inheritance. Our results indicate the importance of modelling the stochasticity and conservation of microbes – only in this way did we appreciate that inheritance can lead to stochastic loss (Fig. 2-3) and that persistence of lineage taxa may be prolonged by asymmetric inheritance (Fig. 5D). Because microbial frequencies are often small, the omission of stochastic effects from models could lead to misestimate the impact of inheritance.

Vertical transfer of microbes might occur in the most diverse host species [12, 18], with only a few exceptions [3]. A great diversity of reproduction and delivery modes might, in turn, determine the distribution of their inheritance – namely the number of microbes transferred and its probability. A comparison of two qualitatively distinct distributions (low and seed-like inheritance in Fig. 1B), indicates they might influence the presence and frequency of microbes differently (Fig. Sup. 5). A consistent cargo in seeds might guarantee the presence of certain microbes in plants [17], who might sometimes benefit from being the first colonizers [28]. In contrast, greater variation might be expected for mammals, where changing amounts of microbes are obtained from epithelia during delivery [11, 12]. Overall, these intrinsic differences might affect the ecological and evolutionary dynamics of hosts and microbes.

We found that microbial inheritance is effective only for some life-histories. While it has been shown that symbiosis [19] and fidelity of inheritance [18] can evolve driven by selection, our results suggest the evolution of life-history traits itself, independent of symbiosis, can impact the relevance of microbial inheritance. Interestingly, the emergence of symbiosis could lead to selection acting on the more evolvable and impactful traits – not only the fidelity of inheritance [18].

Investigating microbial inheritance experimentally poses technical challenges [11]. However, developments using diverse host species [15, 14, 16, 17], suggest that our predictions could be tested experimentally. Firstly, that inheritance is more influential at intermediate host lifespan, environmental migration, or carrying capacity (Fig. 2-3). Related host species with diverse life histories could be compared [32]; alternatively, control could be increased using model organisms amenable to manipulate such traits [33]. Secondly, that the maximum lineage taxa frequency changes with life-history (Eq. 8), could be tested using germ-free or gnotobiotic hosts [17]. Finally, the effect of distinct distributions of microbes inherited (Fig. 1) could be surveyed.

Our approach simplifies the complexity of natural microbiomes. A natural step forward would be considering fitness differences among microbes. These could interact with inheritance to preserve or out-compete certain microbes. Secondly, the host population structure could be included. In such a scenario, subpopulations characterized by different microbiomes could emerge [21]. Moreover, critical connectivity might be needed for inheritance to be effective. Finally, we did not account for specific reproductive ages (or development). This might be particularly relevant because, as we have shown, the effect of inheritance erodes through time.

## 5. Conclusion

Microbial inheritance can influence the occurrence and abundance of microbes within the host-associated commensal microbiome. Even the persistence of microbes absent from the environment could be facilitated in some cases. These findings extend to diverse scenarios of inheritance representative of different host species. However, inheritance is not a silver bullet, instead life-history in terms of environmental immigration, early microbial proliferation, and host lifespan limit its magnitude and temporal extent. Only certain naturally occurring host-microbiome pairs might meet such conditions to exploit its benefits.

## Availability of data

The data generated and analysed during the current study can be simulated from the Python code available via GitHub at https://github.com/romanzapien/microbiome-inheritance.git

## Acknowledgements

We thank the *Evolutionary Theory Department* in the MPI Plön and the *CRC 1182: Origins and Functions of Metaorganisms* for fruitful discussions.

## Authors’ contributions

The original model was developed in discussions between RZC, MS, and AT. RZC analysed the model, programmed the code, and wrote the initial draft. All authors interpreted the results, reviewed the manuscript, and approved the final version.

## A. Appendix

### A. Supplementary methods

#### A.1. Deterministic and stochastic components of the model

We have introduced a model of the microbiome dynamics where we track the frequencies of a taxon *i, x*_*i*_, and the set of other taxa, *o*_*i*_; together, the vector **x** = {*x*_*i*_, *o*_*i*_}. In Eq. (5) we expressed the model in the form of a stochastic differential equation – that describes the microbial dynamics within a host during its lifespan– where the deterministic, **A**[**x**], and stochastic, *B*[**x**], contributions were introduced. Changes have magnitude 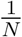. The deterministic part is given by the expected change of **x** that results from the transition probabilities in Eq. (1),

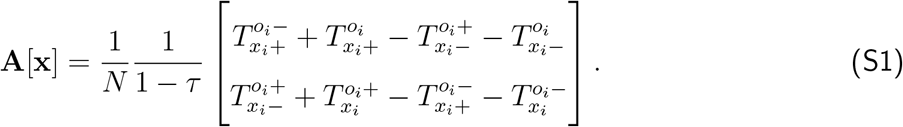

The stochastic part is related to the matrix of covariant change of **x**:

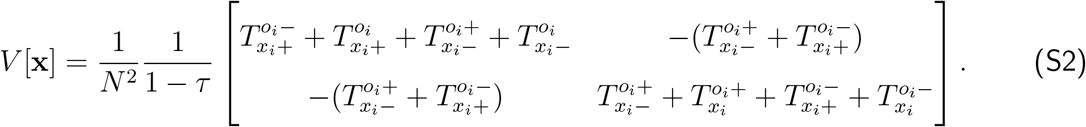

*B*[**x**] is the matrix that satisfies *B*[**x**]^*T*^ *B*[**x**] = *V* [**x**]. This is calculated analytically [34] after defining the quantities 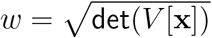 and 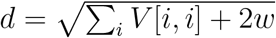

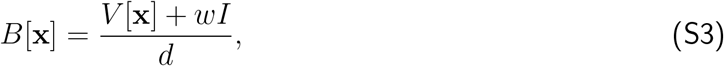

where *I* is the identity matrix.

Note that Eq. (S1) and Eq. (S2) refer to the lifetime of a single host, therefore we divide by 1 − *τ* to remove it from each transition probability. We had introduced 1 − *τ* in Eq. (1) to explain the effect of host death at the population level.

#### A.2. Condition for deterministic increase of lineage taxa

We start from the definition of **A**[1], Eq. (S1). This equation indicates the deterministic change of frequency of a lineage taxon (*x*_*i*_) as a function of *x*_*i*_, other microbes frequency (*o*_*i*_), and parameters of migration (*m*), frequency in the pool of colonizers (*p*_*i*_), and how rapidly available space is occupied (*α*_0_). Asking under which condition **A**[1] > 0, leads to

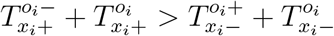

Using the definition of the transition probabilities in Eq. (1) and simplifying, we find

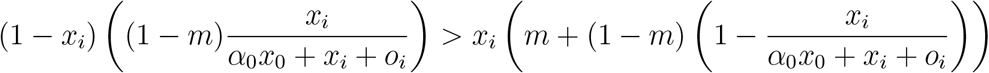

where we used the fact that lineage taxa are absent from the pool of colonizers, *p*_*i*_ = 0. Simplifying and solving for *x*_*i*_ + *o*_*i*_ = 1 − *x*_0_, we find

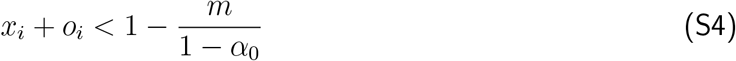

Thus, the growth of lineage taxa stops before the microbial load, *x*_*i*_ + *o*_*i*_, reaches frequency 1, as this is constrained by migration, *m*, and how rapidly available space is occupied, *α*_0_.

### B. Supplementary figures

**Figure Supplementary 1:**
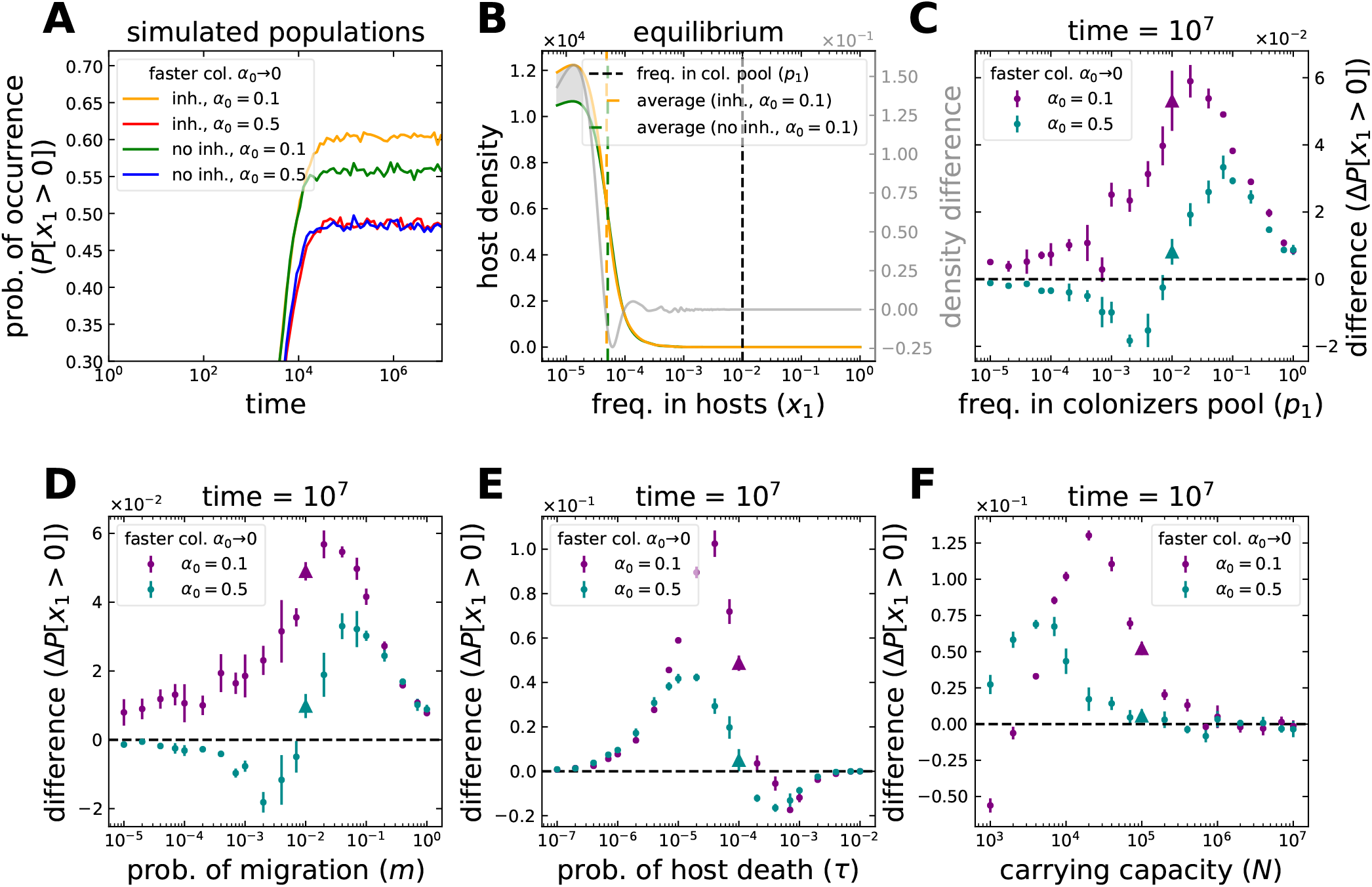
Occurrence of a microbial taxon in hosts under microbial inheritance. We repeat the analysis from Fig. 2, but instead of load, *x*_*i*_ + *o*_*i*_, we look into a single microbial taxon, *x*_*i*_. (**A**) Starting from a condition where all hosts are initially empty, the microbial occurrence increases through time. In this particular case, inheritance increases the occurrence if hosts are colonized rapidly, *α*_0_ → 0. (**B**) The hosts now contain the taxon in small frequencies. The cases shown in (A-B), with parameters *p*_1_ = 10^−2^, *m* = 10^−2^, *τ* = 10^−4^, and *N* = 10^5^, are indicated by the triangles in (C-F). (**C**) Changes are small for other frequencies in the pool of colonizers, *p*_1_, but those at intermediate values benefit the most from inheritance. (**D**) The maximum change occurs for intermediate migration from the pool of colonizers, *m*. For *m* → 1 the taxon colonizes hosts even without inheritance. Instead for *m* → 0 the taxon does not colonize the hosts. (**E**) Larger changes occur for intermediate host death probabilities, *τ*, and fast colonization. Long living hosts, *τ* → 0, contain the taxon even without inheritance. Short living hosts, *τ* → 1, are less likely to be colonized by the taxon within their lifetime. (**F**) In contrast to the microbial load (Fig. 2E), for a single taxon the maximum change occurs at intermediate capacities for microbes, *N*. The change can be negative once inheritance favours more abundant taxa competing for limited space (see C-F). Points and bars in (C-F) indicate the average and standard deviation of 6 simulation pairs, with vs. without inheritance, with 10^4^ hosts each. Offspring receive 9% of their parent’s microbiome on average, *a*_*i*_ = 0 and *b*_*i*_ = 9 in Eq. (4). The whole distributions are shown in Fig. Sup. 3.

**Figure Supplementary 2:**
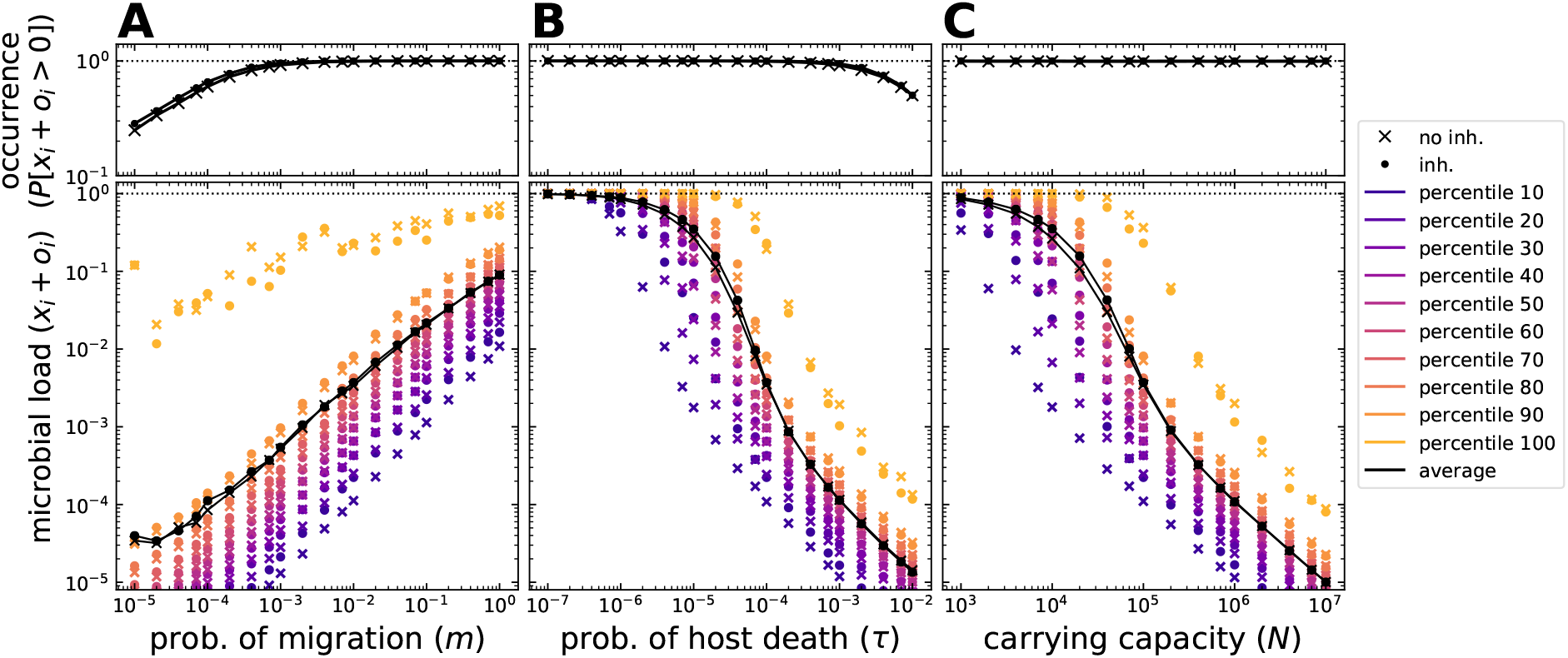
Microbial load distribution across a host population, with or without microbial inheritance. The microbial load is the set of all microbes. In contrast to the difference between distributions, Figs. 2 and 3, here the distributions are shown. The cases without and with inheritance are indicated by × and, •, respectively. Single parameters are modified from the condition *m* = 10^−2^, *τ* = 10^−4^, and *N* = 10^5^. The probability of occurrence and frequencies within hosts increase for (**A**) larger migration from the pool of colonizers, *m* → 1, and (**B**) longer host lifespan, *τ* → 0. (**C**) While occurrence is constant at 1, frequencies increase for smaller capacities for microbes, *N*. Inheritance might increase both observables for certain parameter combinations and percentiles of the distribution (compare • to ×). The increase is evident for small percentiles. Decrease might occur for large percentiles. Only for *τ* ≲ 2 · 10^−7^ all hosts reach carrying capacity within their lifetime. Each simulation included 10^4^ hosts and parameters *a*_*i*_ = 0 and *b*_*i*_ = 9 for inheritance, Eq. (4) – offspring receive 9% of their parent’s microbiome on average – and *α*_0_ = 0.1 for available space occupation.

**Figure Supplementary 3:**
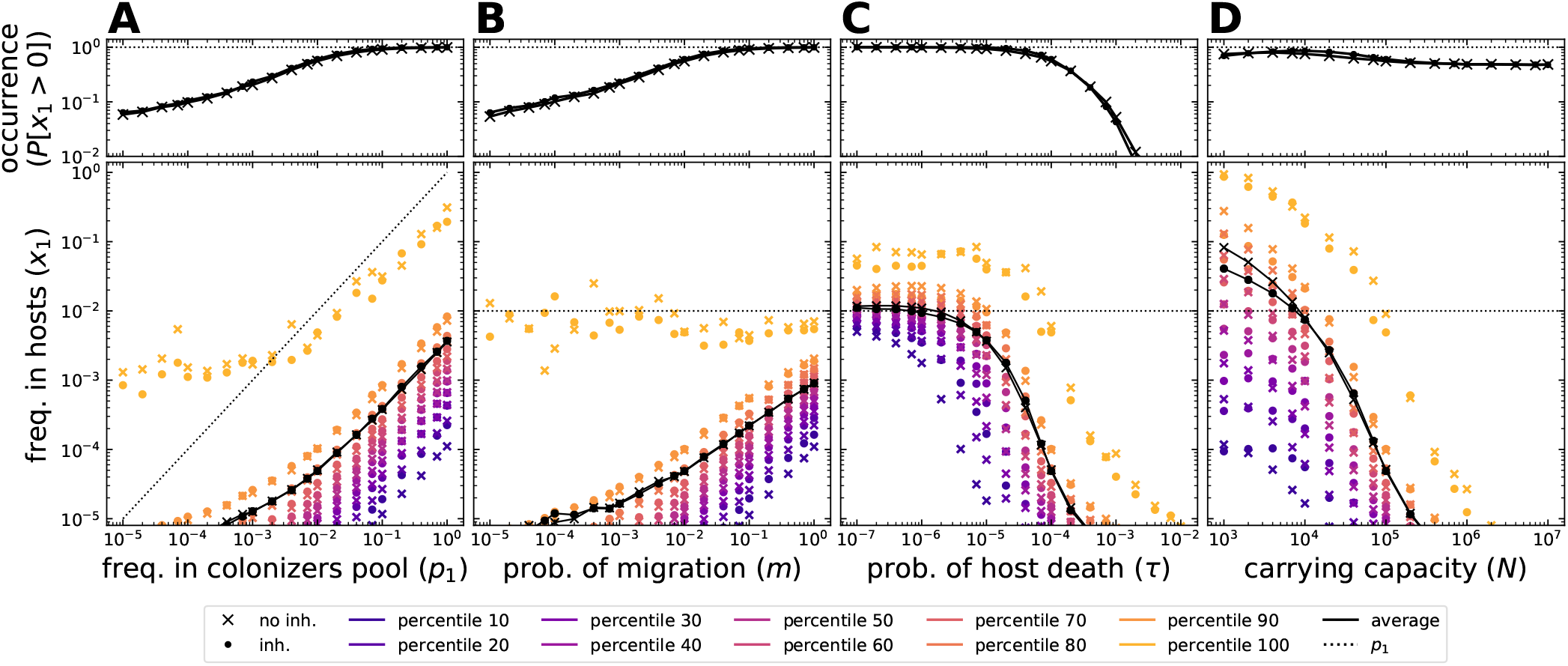
Frequency of a microbial taxon distribution across the host population, with or without inheritance. In contrast to the difference between distributions, Figs. Sup. 1 and 4, here the distributions are shown. The cases without and with inheritance are indicated by × and, •, respectively. Single parameters are modified from the condition *p*_1_ = 10^−2^, *m* = 10^−2^, *τ* = 10^−4^, and *N* = 10^5^. (**A**) The probability of occurrence and frequency within hosts increase for higher abundances in the pool of colonizers, *p*_1_ → 1, and (**B**) larger migration from the environment, *m* → 1. For *p*_1_ → 0, hosts with larger frequencies than in the pool of colonizers (*x*_1_ *> p*_1_) might occur stochastically. In contrast to microbial load (Fig. Sup. 2), inheritance might decrease the frequencies for (**C**) long host lifespans, *τ* → 0, and, (**D**) smaller capacities for microbes, *N*, where hosts are fully colonized. The reduced variability of the early microbiome, makes hosts with initially large frequencies of the microbial taxon less likely. Even if low frequencies increase, the average frequency decreases as a result. Inheritance increases the average frequency for intermediate values of *τ* and *N*, where hosts are partially colonized (Fig. Sup. 2 B-C). Each simulation included 10^4^ hosts and parameters *a*_*i*_ = 0 and *b*_*i*_ = 9 for inheritance, Eq. (4) – offspring receive 9% of their parent’s microbiome on average – and *α*_0_ = 0.1 for the available space occupation.

**Figure Supplementary 4:**
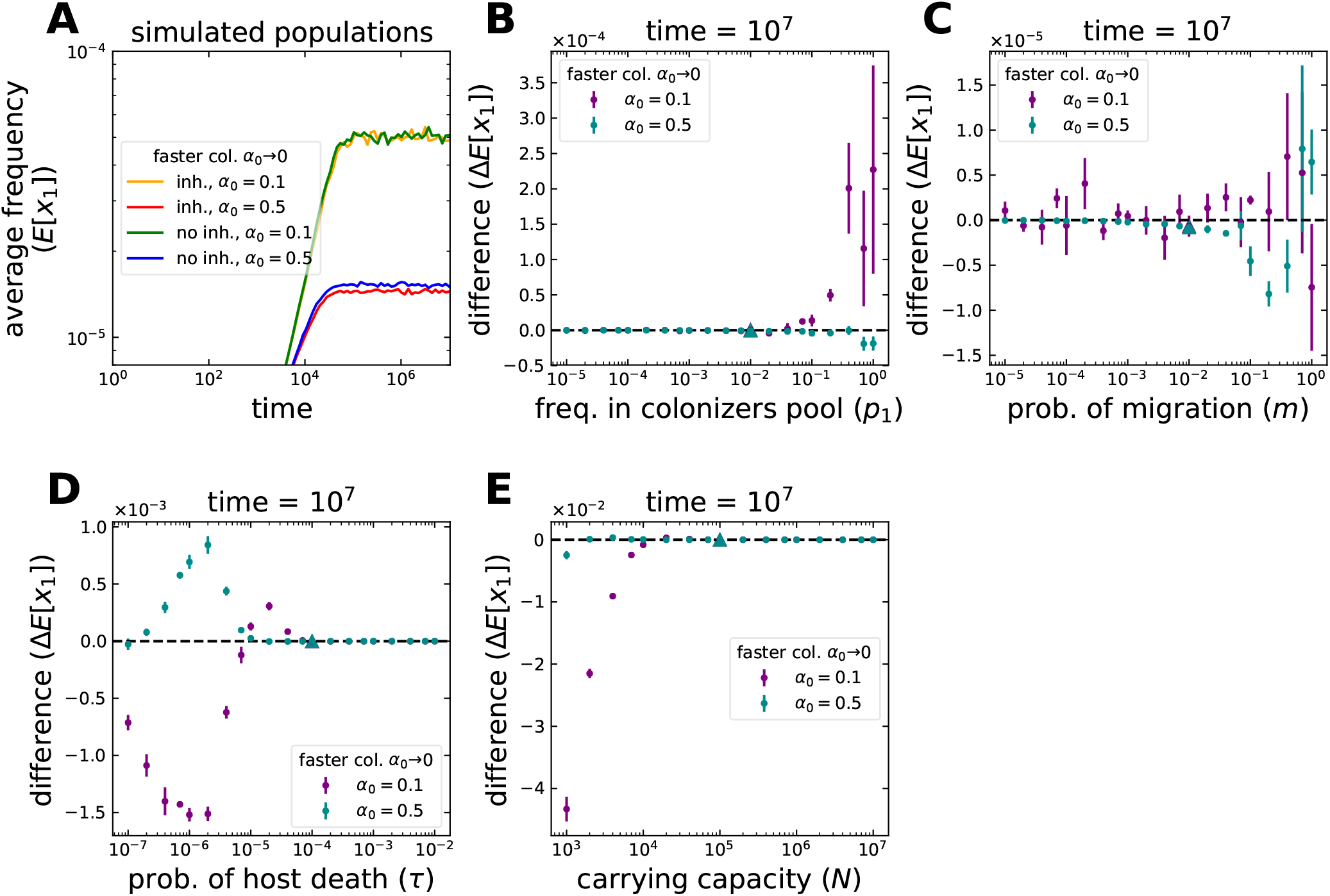
Average frequency of a microbial taxon in hosts under microbial inheritance. We repeat the analysis from Fig. 3, but instead of load, *x*_*i*_ + *o*_*i*_, we look into a single microbial taxon, *x*_*i*_. (**A**) Starting from a condition where all hosts are initially empty, the average frequency of microbes in hosts increases through time before reaching an equilibrium. In this particular case, inheritance makes the average slightly larger if hosts are occupied more slowly, *α*_0_ = 0.5. Although more hosts harbour the taxon, no change occurs for *α*_0_ = 0.1, as inheritance reduces the variability between individuals. The cases shown in (A), with parameters *p*_1_ = 10^−2^, *m* = 10^−2^, *τ* = 10^−4^, and *N* = 10^5^, are indicated by the triangles in (B-E). (**B**) No changes occur for multiple frequencies in the pool of colonizers, *p*_1_, and (**C**) migrations from the pool of colonizers, *m*. (**D**) The largest changes occur for intermediate host death probabilities, *τ*. For long living hosts, *τ* → 0, the change produced by inheritance can be negative. (**E**) Similarly for small capacities for microbes, *N*, where inheritance causes abundant taxa to outcompete others. Points and bars in (B-E) indicate the average and standard deviation of 6 simulation pairs, with vs. without inheritance, with 10^4^ hosts each. Offspring receive 9% of their parent’s microbiome on average, *a*_*i*_ = 0 and *b*_*i*_ = 9 in Eq. (4). The whole distributions are shown in Fig. Sup. 3.

**Figure Supplementary 5:**
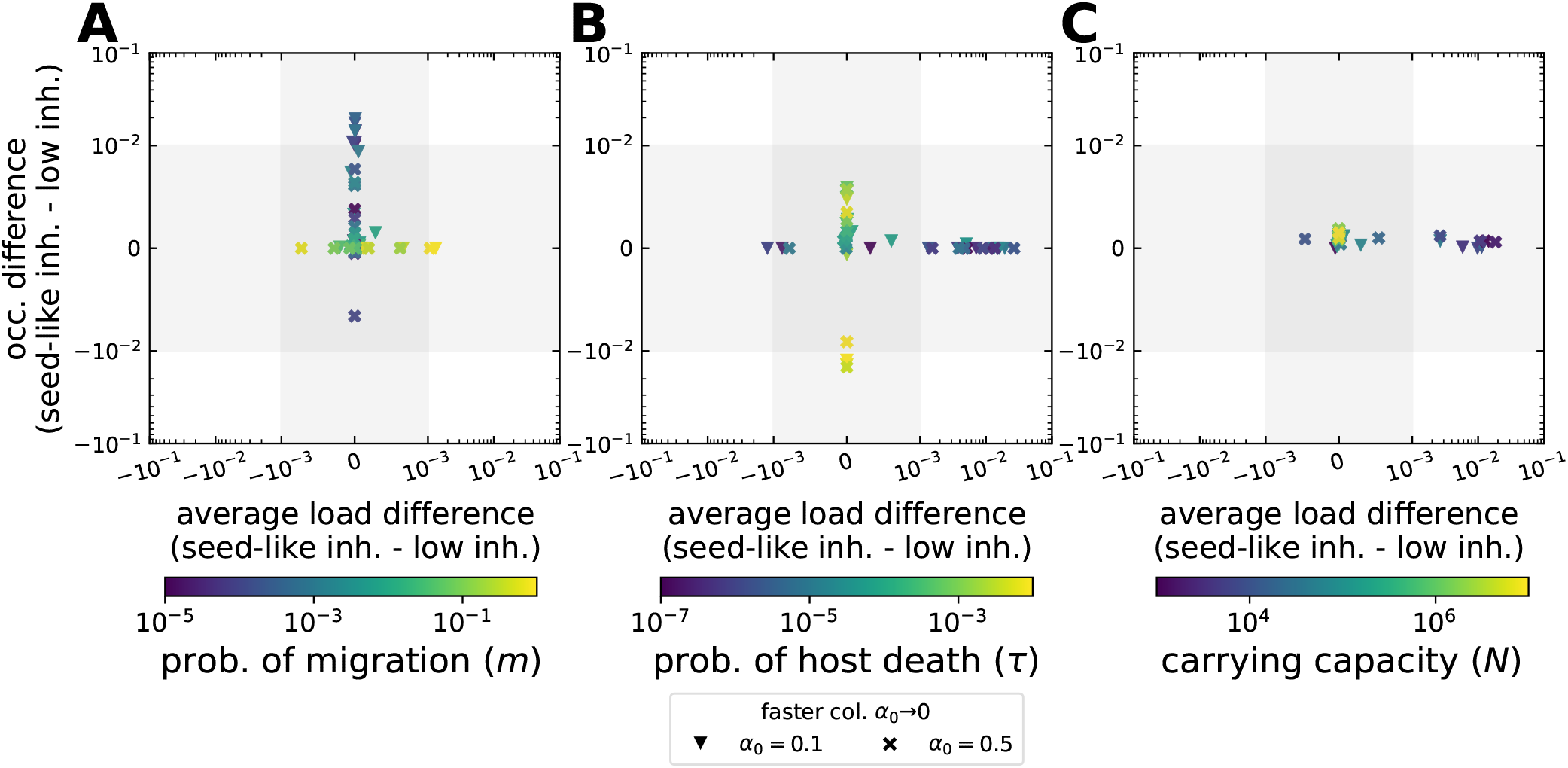
Difference in microbial load between ‘low’ and ‘seed-like’ inheritance. A positive difference indicates the observable is larger for seed-like inheritance (Fig. 1B). For both, low and seed-like inheritance, offspring receive 9% of their parent’s microbiome on average (*a*_*i*_ = 0 and *b*_*i*_ = 9 for low inheritance, and *a*_*i*_ = 9 and *b*_*i*_ = 99 for seed-like inheritance in Eq. (4)). Low inheritance corresponds to data shown in Fig. 2 and Fig. 3. Single parameters are modified from the condition *m* = 10^−2^, *τ* = 10^−4^, and *N* = 10^5^. (**A**) For low migration from the pool colonizers, *m* → 0, seed-like inheritance increases the microbial occurrence (a exception stems from a slower occupation of available space, *α*_0_ = 0.5). For *m* → 1, it mildly increases the average microbial load. (**B**) For low host death, *τ* → 0, this inheritance mode increases the average load importantly. For *τ* → 1, it only affects the occurrence, even decreasing it. (**C**) For varying carrying capacity (*N*), larger average loads are obtained for small *N*. Each point corresponds to the difference of observables calculated from simulations with 10^4^ hosts. The scale of axes is logarithmic, but linear within [−10^−3^, 10^−3^] for the average load, and [−10^−2^, 10^−2^] for the occurrence.

**Figure Supplementary 6:**
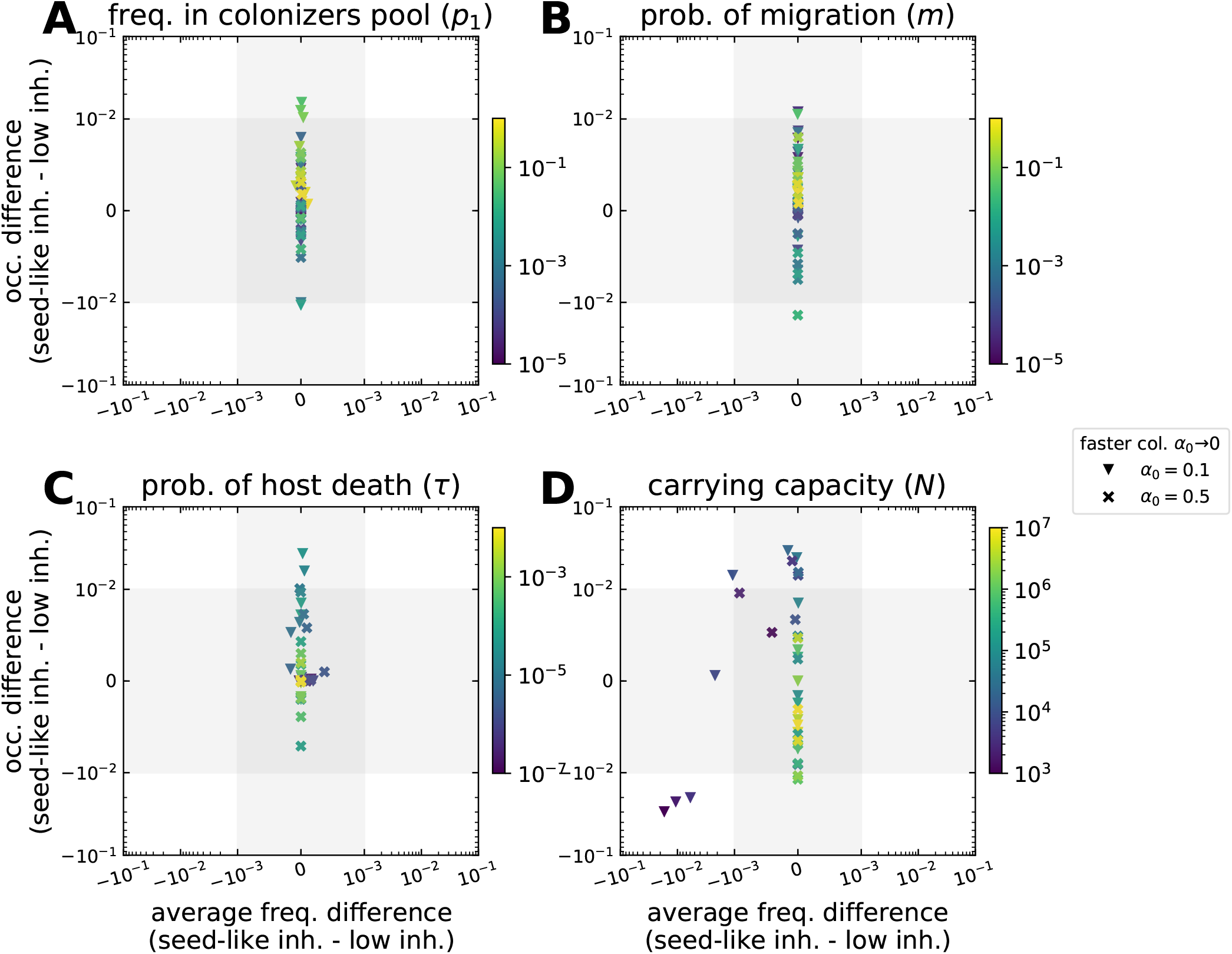
Difference in the frequency of a microbial taxon between ‘low’ and ‘seed-like’ inheritance. A positive difference indicates the observable is larger for seed-like inheritance (Fig. 1B). For both, low and seed-like inheritance, offspring receive 9% of their parent’s microbiome on average (*a*_*i*_ = 0 and *b*_*i*_ = 9 for low inheritance, and *a*_*i*_ = 9 and *b*_*i*_ = 99 for seed-like inheritance in Eq. (4)). Low inheritance corresponds to data shown in Fig. Sup. 1 and Fig. Sup. 4. Single parameters are modified from the condition *p*_1_ = 10^−2^, *m* = 10^−2^, *τ* = 10^−4^, and *N* = 10^5^. (**A**-**C**) A seed-like inheritance primarily modifies the occurrence for various values of frequency in the pool of colonizers (*p*_*i*_), migration (*m*), and host death (*τ*). (**D**) For varying values of the carrying capacity for microbes (*N*), the main change is on the occurrence, however, for small *N* a decrease of average frequency is observed. A decrease or increase of occurrence is not clearly attributable to the rate of host colonization (*α*_0_). Each point corresponds to the difference of simulations with 10^4^ hosts. The scale of axes is logarithmic, but linear within [−10^−3^, 10^−3^] for the average frequency, and [−10^−2^, 10^−2^] for the occurrence.

## References

[1] Luke R Thompson, Jon G Sanders, Daniel McDonald, Amnon Amir, Joshua Ladau, Kenneth J Locey, Robert J Prill, Anupriya Tripathi, Sean M Gibbons, Gail Ackermann, et al. A communal catalogue reveals earth’s multiscale microbial diversity. Nature, 551(7681):457–463, 2017.

[2] Timothy J Colston and Colin R Jackson. Microbiome evolution along divergent branches of the vertebrate tree of life: what is known and unknown. Molecular Ecology, 25(16):3776–3800, 2016.

[3] Tobin J Hammer, Jon G Sanders, and Noah Fierer. Not all animals need a microbiome. FEMS Microbiology Letters, 366(10):fnz117, 2019.

[4] Julia Johnke, Philipp Dirksen, and Hinrich Schulenburg. Community assembly of the native c. elegans microbiome is influenced by time, substrate and individual bacterial taxa. Environmental Microbiology, 22(4):1265–1279, 2020.

[5] Elizabeth T. Miller and Brendan J. M. Bohannan. Life Between Patches: Incorporating Mi-crobiome Biology Alters the Predictions of Metacommunity Models. Frontiers in Ecology and Evolution, 7, 2019.

[6] M. Sieber, A. Traulsen, H. Schulenburg, and A. E. Douglas. On the evolutionary origins of host-microbe associations. Proceeding of the National Academy of Sciences, 118(9):e2016487118, 2021.

[7] Alexandre Almeida, Alex L Mitchell, Miguel Boland, Samuel C Forster, Gregory B Gloor, Aleksandra Tarkowska, Trevor D Lawley, and Robert D Finn. A new genomic blueprint of the human gut microbiota. Nature, 568(7753):499–504, 2019.

[8] Braedon McDonald and Kathy D McCoy. Maternal microbiota in pregnancy and early life. Science, 365(6457):984–985, 2019.

[9] Monika Bright and Silvia Bulgheresi. A complex journey: transmission of microbial symbionts. Nature Reviews Microbiology, 8(3):218–230, 2010.

[10] Raphael Eisenhofer, Jeremiah J Minich, Clarisse Marotz, Alan Cooper, Rob Knight, and Laura S Weyrich. Contamination in low microbial biomass microbiome studies: issues and recommendations. Trends in microbiology, 27(2):105–117, 2019.

[11] Maria Elisa Perez-Muñoz, Marie-Claire Arrieta, Amanda E Ramer-Tait, and Jens Walter. A critical assessment of the “sterile womb” and “in utero colonization” hypotheses: implications for research on the pioneer infant microbiome. Microbiome, 5(1):48, 2017.

[12] Lisa J Funkhouser and Seth R Bordenstein. Mom knows best: the universality of maternal microbial transmission. PLoS Biol, 11(8):e1001631, 2013.

[13] Shelbi L Russell. Transmission mode is associated with environment type and taxa across bacteria-eukaryote symbioses: a systematic review and meta-analysis. FEMS microbiology letters, 366(3):fnz013, 2019.

[14] Johannes R Björk, Cristina Díez-Vives, Carmen Astudillo-García, Elizabeth A Archie, and José M Montoya. Vertical transmission of sponge microbiota is inconsistent and unfaithful. Nature Ecology & Evolution, page 1, 2019.

[15] Andrew H Moeller, Taichi A Suzuki, Megan Phifer-Rixey, and Michael W Nachman. Transmission modes of the mammalian gut microbiota. Science, 362(6413):453–457, 2018.

[16] Justinn Renelies-Hamilton, Kristjan Germer, David Sillam-Dussés, Kasun H Bodawatta, and Michael Poulsen. Disentangling the relative roles of vertical transmission, subsequent colonizations, and diet on cockroach microbiome assembly. Msphere, 6(1), 2021.

[17] Ezgi Özkurt, M Amine Hassani, Uğur Sesiz, Sven Künzel, Tal Dagan, Hakan Özkan, and Eva H Stukenbrock. Higher stochasticity of microbiota composition in seedlings of domesticated wheat compared to wild wheat. bioRxiv, page 685164, 2019.

[18] Marjolein Bruijning, Lucas P Henry, Simon KG Forsberg, C Jessica E Metcalf, and Julien F Ayroles. When the microbiome defines the host phenotype: selection on vertical transmission in varying environments. bioRxiv, 2020.

[19] Simon Van Vliet and Michael Doebeli. The role of multilevel selection in host microbiome evolution. PNAS, 116(41):20591–20597, 2019.

[20] Joan Roughgarden. Holobiont evolution: Mathematical model with vertical vs. horizontal microbiome transmission. Philosophy, Theory, and Practice in Biology, 12(2), 2020.

[21] Philip T Leftwich, Matthew P Edgington, and Tracey Chapman. Transmission efficiency drives host–microbe associations. Proceedings of the Royal Society B, 287(1934):20200820, 2020.

[22] Shen Jean Lim and Seth R Bordenstein. An introduction to phylosymbiosis. Proceedings of the Royal Society B, 287(1922):20192900, 2020.

[23] Qinglong Zeng, Jeet Sukumaran, Steven Wu, and Allen Rodrigo. Neutral models of microbiome evolution. PLoS Computational Biology, 11(7):e1004365, 2015.

[24] Fan Zhang, Maureen Berg, Katja Dierking, Marie-Anne Félix, Michael Shapira, Buck S Samuel, and Hinrich Schulenburg. Caenorhabditis elegans as a model for microbiome research. Frontiers in Microbiology, 8:485, 2017.

[25] Jessamina E Blum, Caleb N Fischer, Jessica Miles, and Jo Handelsman. Frequent replenishment sustains the beneficial microbiome of drosophila melanogaster. MBio, 4(6), 2013.

[26] Juan M Cancino and NH Rodger. An ecological overview of cloning in metazoa. Population biology and evolution of clonal organisms, pages 153–186, 1985.

[27] Wolfgang Frey and Harald Kürschner. Asexual reproduction, habitat colonization and habitat maintenance in bryophytes. Flora, 206(3):173–184, 2011.

[28] R. Zapien-Campos, M. Sieber, and A. Traulsen. Stochastic colonization of hosts with a finite lifespan can drive individual host microbes out of equilibrium. PLoS Computational Biology, 16(11):e1008392, 2020.

[29] I. S. Gradshteyn and I. M. Ryzhik. Table of Integrals, Series and Products. Academic Press, London, 1994.

[30] C. W. Gardiner. Handbook of Stochastic Methods. Springer, NY, third edition, 2004.

[31] Hilary P. Browne, Samuel C. Forster, Blessing O. Anonye, Nitin Kumar, B. Anne Neville, Mark D. Stares, David Goulding, and Trevor D. Lawley. Culturing of ‘unculturable’ human microbiota reveals novel taxa and extensive sporulation. Nature, 533(7604):543–546, May 2016.

[32] Se Jin Song, Jon G Sanders, Frédéric Delsuc, Jessica Metcalf, Katherine Amato, Michael W Taylor, Florent Mazel, Holly L Lutz, Kevin Winker, Gary R Graves, et al. Comparative analyses of vertebrate gut microbiomes reveal convergence between birds and bats. MBio, 11(1), 2020.

[33] Thomas CG Bosch, Karen Guillemin, and Margaret McFall-Ngai. Evolutionary “experiments” in symbiosis: The study of model animals provides insights into the mechanisms underlying the diversity of host–microbe interactions. BioEssays, 41(10):1800256, 2019.

[34] Edward Allen. Modeling with Itô Stochastic Differential Equations. Springer, 2007.

